# Accurate somatic variant calling performance using Avidity Sequencing with Burning Rock’s OncoScreen^TM^ Plus panel

**DOI:** 10.1101/2023.09.20.558622

**Authors:** Guangliang Zhang, Yangyang Liu, Yilun Luo, Yusheng Han, Zhihong Zhang

## Abstract

**Background:** Next-generation sequencing (NGS) comprehensive genomic profiling panels provide targeted genome-wide detection of the somatic variant landscape of various cancer types. Recently, an innovative sequencing technology, Avidity Sequencing from Element Biosciences, has emerged to provide economical option for mid-throughput sequencing, with high quality (Q40).

**Aims:** The aim of this study was to evaluate the performance characteristics of AVITI sequencing by avidity on detection of somatic variants from reference samples, and to compare the variant detection concordance between AVITI and NovaSeq sequencing platforms by evaluating the variant callings of 518 cancer-related genes using Burning Rock’s OncoScreen^TM^ Plus panel.

**Methods:** Contrived commercial reference control samples (harboring known variants) and clinical Formalin-Fixed Paraffin-Embedded tissue (FFPE) samples were examined to evaluate the variant detection performance of two sequencing platforms, with respects to Single Nucleotide Variant (SNV), Insertion-Deletion (Indel), Structure Variant (SV/Fusion), Copy Number Variation (CNV), Microsatellite Instability (MSI), and Tumor Mutation Burden (TMB). Variant specific QC metrics were developed to evaluate sequencing-level and variant calling level accuracy. All samples were processed utilizing the OncoScreen^TM^ Plus assay, including library preparation and bioinformatics post sequencing data analysis.

**Results:** The OncoScreen^TM^ Plus assay is compatible with the AVITI system. Samples prepared with the OncoScreen^TM^ Plus panel and sequenced on the AVITI system has successfully detected cancer related variants including SNV/Indels, SV/Fusion, CNVs, MSIs, and TMB, which were highly concordant with reference controls. In addition, the AVITI system produced a higher overall quality score, index assignment rate, mean target coverage, and lower optical duplication rate AVITI than NovaSeq system.

**Conclusion:** The new sequencing technology Avidity from AVITI system can be applied seamlessly to current high-performance targeted oncology assays to reliably identify somatic variants in a flexible and cost-effective manner.

## Introduction

In the recent decade, morbidity and mortality of cancer has been significantly decreased, aided in part by NGS based Comprehensive Genomic Profiling (CGP) panel, which were designed to detect multiple types of somatic variants in cancer specimens while achieving high sensitivity and close to 100% specificity. CGP assays have a broad range of clinical applications, such as identification of actionable variant classes: Single Nucleotide Variants (SNV), Insertion-Deletions (InDel), Structure Variants (SV/Fusion), and Copy Number Variants (CNV), as well as exploration of drug resistance, and assessment of immunotherapy related genomic signatures such as micro-satellite instability (MSI) and Tumor Mutation Burden(TMB) (1–5). As a complex application of NGS technology, CGP panels pose increasingly high demands for sequencing quality and accuracy in order to improve data analysis sensitivity and specificity.

OncoScreen^TM^ Plus Cancer Mutation Profiling Tissue Kit is designed to increase the coverage of actionable variants and to improve the assay sensitivity. This kit is intended to be used for the qualitative detection of multiple types of variants from patient FFPE tissue samples with various cancers involving 518 gene targets, including SNV, Indel, Structure Variants (SV/Fusion), CNV, as well as the determination of microsatellite instability (MSI) status and Tumor Mutation Burden (TMB). This assay has previously been used to successfully detect variants in early onset metastatic colorectal cancer samples using Illumina’s sequencing instruments (6). While Illumina sequencers have become the most widely adopted platform, the high instrument and reagent cost hinders broader applications (7). Recently, several new NGS technologies have entered the market. One example is sequencing by Avidity from Element Biosciences (8). A comprehensive study to compare newly released technologies for performance quality and concordance is necessary in order to drive adoption of new sequencing technology and advance future oncology research. Moreover, this continued re-assessment is of particular importance as researchers implement new sequencing technologies in a clinical setting (9, 10).

In this study, we evaluated a novel sequencing technology, Sequencing by Avidity. Avidity Sequencing separates and optimizes, respectively, the process of traversing through a DNA template and the process of identifying each nucleotide within the template, which achieves accuracy surpassing Q40, enabling a diversity of applications. We compared the variant calling performance and concordance between the AVITI and NovaSeq platforms across 518 cancer-related genes based on OncoScreen^TM^ Plus panel. In order to evaluate different biomarker detected in this CGP panel, we developed variant-specific QC metrics for both sequencing-level and secondary analysis level. This study generated comparable results between sequencing instruments while also discovered some platform-specific features. Using the aforementioned parameters we were able to confirm a list of true variants within the contrived DNA reference control samples tested in this study, which can serve as a reference data set of reliable and consistent control variants for future oncology study and technical comparison.

## Materials and methods

### Study design

In Study 1 and Study 2, contrived control reference samples (harboring known variants) and clinical FFPE tissue samples were used. Detailed information of these samples is listed in Table 1. In Study 1, contrived tumor samples of HD734 and HD833 were purchased from Horizon Discovery. NA19240 was purchased from the Coriell Institute. HD833 was diluted to 1/2, 1/4, 1/8, and 1/16 using NA19240 based on DNA quantity measured by Qubit (Thermo Fisher). In Study 2, 8 FFPE tumor samples were purchased from Discovery Life Science and 6 reference control samples (HD728, HD729, HD730, HD731, HD753, and HD827) were purchased from Horizon Discovery. In addition, 2 internal reference control samples were prepared by Burning Rock Dx. All samples were prepared into NGS libraries with OncoScreen^TM^ Plus assay per standard library preparation methods. Final libraries were sequenced on both the NovaSeq 6000 and Element AVITI systems following vendor manuals. The complete study design is shown in Figure 1.

**Figure 1.**
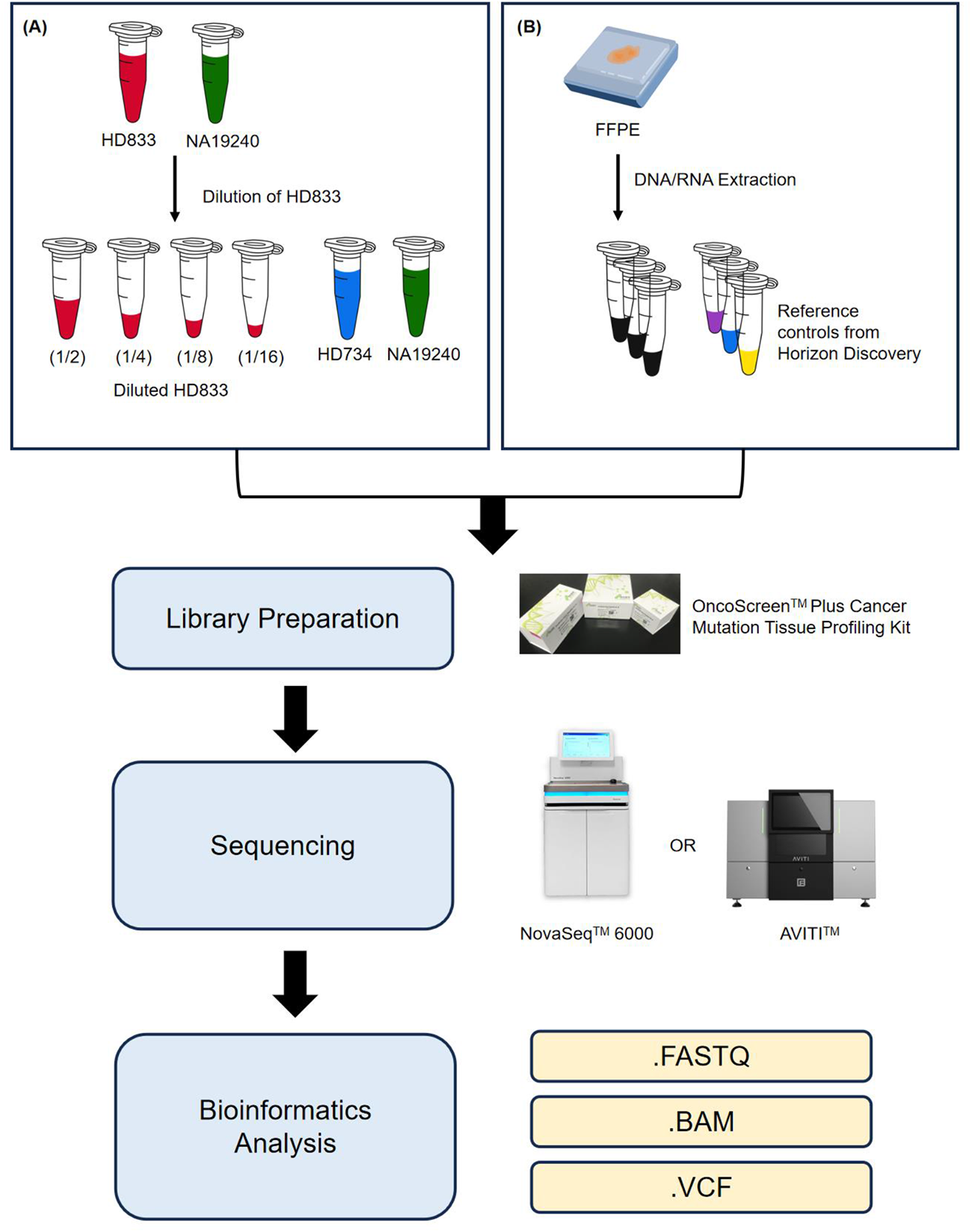
The flowchart of combinations using two sequencing platforms A: serial diluted and original contrived samples were used to evaluate the performance of SNV/Indel detection of OncoScreen^TM^ Plus on both sequencing platforms. B: contrived samples and clinical FFPE samples was used to evaluate the performance of SNV, Indel, Fusion, and CNV detection of OncoScreen^TM^ Plus on both sequencing platforms.

**Table 1.** Information for the samples enrolled in the study.

| **Study** | **Sample Name** | **Sample (Tumor) Type** | **Vendor** |
| --- | --- | --- | --- |
| Study 1 | NA19240 | Cell Line | Coriell Institute |
|  | HD734 | Commercial Control | Horizon Discovery |
|  | HD833 (1/2) | Commercial Control | Horizon Discovery |
|  | HD833 (1/4) | Commercial Control | Horizon Discovery |
|  | HD833 (1/8) | Commercial Control | Horizon Discovery |
|  | HD833 (1/16) | Commercial Control | Horizon Discovery |
| Study 2 | F00117404 | FFPE (Colorectal) | Discovery Life Science |
|  | F00123483 | FFPE (Colorectal) | Discovery Life Science |
|  | F00121163 | FFPE (Gastric) | Discovery Life Science |
|  | F00123076 | FFPE (Pancreatic) | Discovery Life Science |
|  | F00114726 | FFPE (Head and Neck) | Discovery Life Science |
|  | F00115833 | FFPE (Endometrial) | Discovery Life Science |
|  | F00115556 | FFPE (Endometrial) | Discovery Life Science |
|  | F00114353 | FFPE (Endometrial) | Discovery Life Science |
|  | HD827 | Commercial Control | Horizon Discovery |
|  | HD753 | Commercial Control | Horizon Discovery |
|  | HD728 | Commercial Control | Horizon Discovery |
|  | HD729 | Commercial Control | Horizon Discovery |
|  | HD730 | Commercial Control | Horizon Discovery |
|  | HD731 | Commercial Control | Horizon Discovery |

FFPE: paraffin-embedded tissue.

### Library preparation

The design of OncoScreen^TM^ Plus assay is based on an RNA probe capture technique. 100ng of DNAs extracted from control samples, FFPE, or white blood cell (WBC) samples went through DNA fragment shearing, end repair, adapter ligation, purification, and then amplified by polymerase chain reaction (PCR) to prepare a whole-genome pre-library. Amplified DNA was then hybridized with RNA probes of specific sequence to specifically capture all or part of exons regions as well as some important intron regions from 518 genes in the human genome. In total, 22 prepared samples were prepared in replicates in study 1 and in single in study 2 with OncoScreen^TM^ Plus according to the user manual. DNA pre-library fragments captured by the probe were further purified with streptavidin beads and amplified to obtain the final library for sequencing.

### Sequencing with NovaSeq and AVITI

The prepared libraries were pooled and sequenced on NovaSeq^TM^ 6000 System using 2x150 read configuration with index reads following vendor’s manual. Similarly, on the AVITI system, 0.5 pmol (30 µl of 16.7nM) of the same libraries in Study 1 and Study 2 were processed respectively using Adept Compatibility Workflow Kit (Element Biosciences, Cat# 830-00003). Final circularized libraries for Study 1 and Study 2 were quantified using quantitative polymerase chain reaction (qPCR) standard with primer mix provided in the Adept Compatibility Workflow Kit. The quantified library was denatured and sequenced on Element AVITI system using 2x150 read with indexing. FASTQ files generated from AVITI system were used for downstream data analysis.

### Data analysis

Output sequencing data (FASTQ files) from each sequencing systems were evaluated through bioinformatics QC and analyzed using OncoScreen^TM^ Plus Bioinformatics Analysis Pipeline developed by Burning Rock Dx. The analysis software encompasses the entire process from FASTQ data input to report generation output. In the analysis software, the data preprocessing module [based on the Trimmomatic software (11)] was used to remove adapter sequences introduced during library preparation and low-quality base fragments. The sequence alignment module (based on BWA software) was used to align the base sequence in the FASTQ file to the hg19 (GRCh37) human reference genome to generate a BAM file and carry out sorting on the BAM file according to genome coordinates (based on the Picard software). The variant calling module (based on the VarDict software) was used to analyze SNV and Indel variants in the sample. The integration analysis module (developed in-house) was used to analyze structure variants (SV/fusions) in the sample. The CNV analysis module (developed in-house) was used to analyze CNVs in the sample. CNV binning is used to calculate the sequencing depth uniformity of each sample through analyzing the fluctuation of each bin’s coverage. This metric reflects the coverage precision across the entire bed file. The MSI detection module (developed in-house) was used to calculate an MSI score and assess potential MSIs in the sample. MSI score is grouped based on a cut-off to MSI-H or microsatellite stable (MSS). The TMB detection module (developed in-house) was used to analyze TMB score in the sample.

## Results

### Compatibility of the OncoScreen^TM^ Plus assays with the AVITI system

The OncoScreen^TM^ plus assay was originally designed to generate NGS libraries and sequence on Illumina platforms. In order to sequence on Element’s AVITI system, an Adept Compatibility workflow from Element Biosciences was used to convert the final library into a compatible format (Figure 2). The final libraries were quantified, pooled, and sequenced on NovaSeq 6000 or processed through the Adept Compatibility workflow for sequencing on the AVITI system. The sequencing length for both sequencers was 151-bp paired end with 8-bp unique dual indices (UDIs). The FASTQ files generated by the AVITI system were used as input without additional processing for OncoScreen^TM^ Plus Bioinformatics Analysis Pipeline, which was initially developed and optimized based on sequencing data generated from Illumina sequencer by Burning Rock Dx.

**Figure 2.**
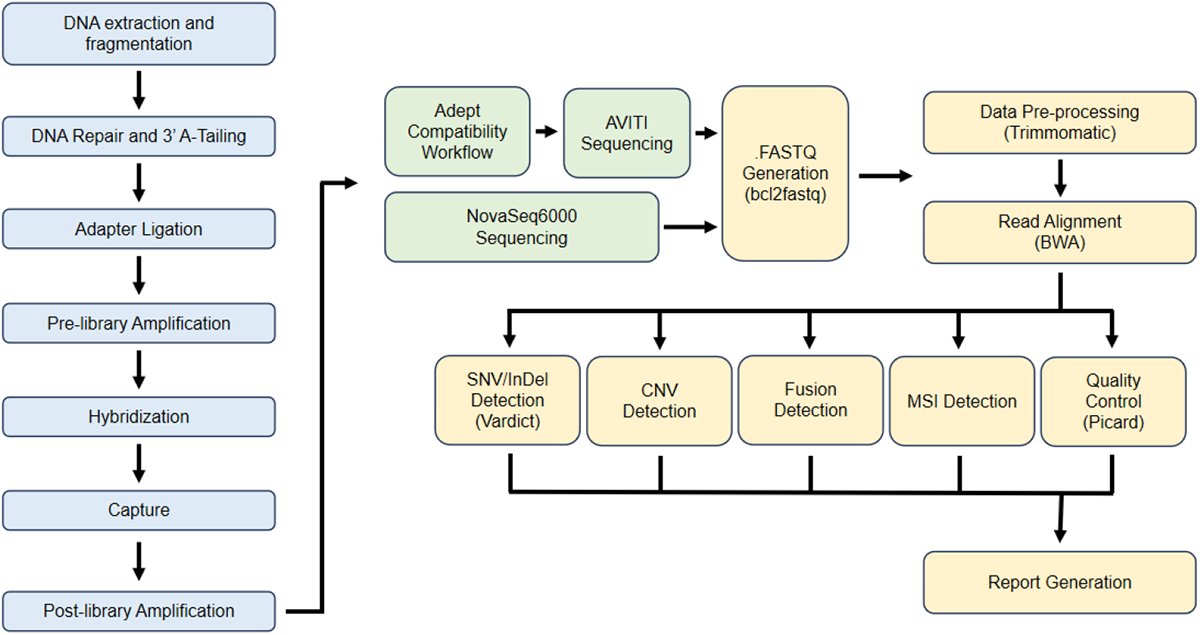
The workflow of Burning Rock OncoScreen^TM^ Plus in library preparation, sequencing on Element AVITI, and analysis pipeline

### Quality score and index assignment

Samples in Study 1 were sequenced twice on the AVITI system (AVITI run 1 and run 2) and once on NovaSeq 6000 SP flow cell (NovaSeq run1). Total output data size from the NovaSeq run in Study 1 was 323Gb, while AVITI runs generated 266Gb and 237Gb in run 1, and 296Gb in run2, comparable to the 286Gb output generated from the NovaSeq SP flow cell (Table 2). The AVITI platform generated a higher percentage of bases with Phred score of Q30 or greater (94% in study 1, 95% in study 2) vs NovaSeq (about 91% in study 1, 89% in study 2). Data from AVITI data also showed a good percentage of bases above Phred quality Q40 (78.05% in run 1 and 82.25% in run 2 in Study 1, and 82% in Study 2). A higher index assignment percentage was obtained from AVITI data (∼99%) compared to NovaSeq (∼93%). All runs on both platforms generated enough high-quality reads for downstream variant calling analysis.

**Table 2.** Sequencing quality metrics.

| **Study** | **Run ID** | **Total yield (Gb)** | **Total reads (M)** | **Q30 (%)** | **Read configuration (bp)** | **Index assigned (%)** |
| --- | --- | --- | --- | --- | --- | --- |
| Study 1 | NOVA-Run1 | 322.99 | 1,029 | 90.8 | 150+150 | 93.4 |
|  | AVITI-Run1 | 266.10 | 897 | 93.8 | 150+150 | 98.9 |
|  | AVITI-Run2 | 236.67 | 905 | 94.9 | 150+150 | 99.3 |
| Study 2 | NOVA-Run2 | 286.53 | 913 | 88.6 | 150+150 | 89.2 |
|  | AVITI-Run3 | 295.96 | 1,001 | 95.1 | 150+150 | 98.5 |

### Optical Duplicate analysis

FASTQ files generated on both platforms of each OncoScreen^TM^ Plus library were aligned against the human reference genome. The resulting raw BAM file was used to evaluate duplicates level using SAMtools (http://enseqlopedia.com/2016/05/increased-read-duplication-on-patterned-flowcells-understanding-the-impact-of-exclusion-amplification/). Duplicates were flagged by Picard MakeDuplicates analysis. Significantly lower optical duplication rates were observed in AVITI data (< 0.45%) compared to NovaSeq data (∼ 8%) in Study 1, and a similar trend was observed in Study 2 (AVITI < 0.2%, NovaSeq < 6.2%) (Figure 3). This result highlights differences in AVITI’s polony amplification technology compared to the EXAmp amplification method and the patterned flow cell used on NovaSeq (https://www.illumina.com/science/technology/next-generation-sequencing/sequencing-technology/patterned-flow-cells.html).

**Figure 3.**
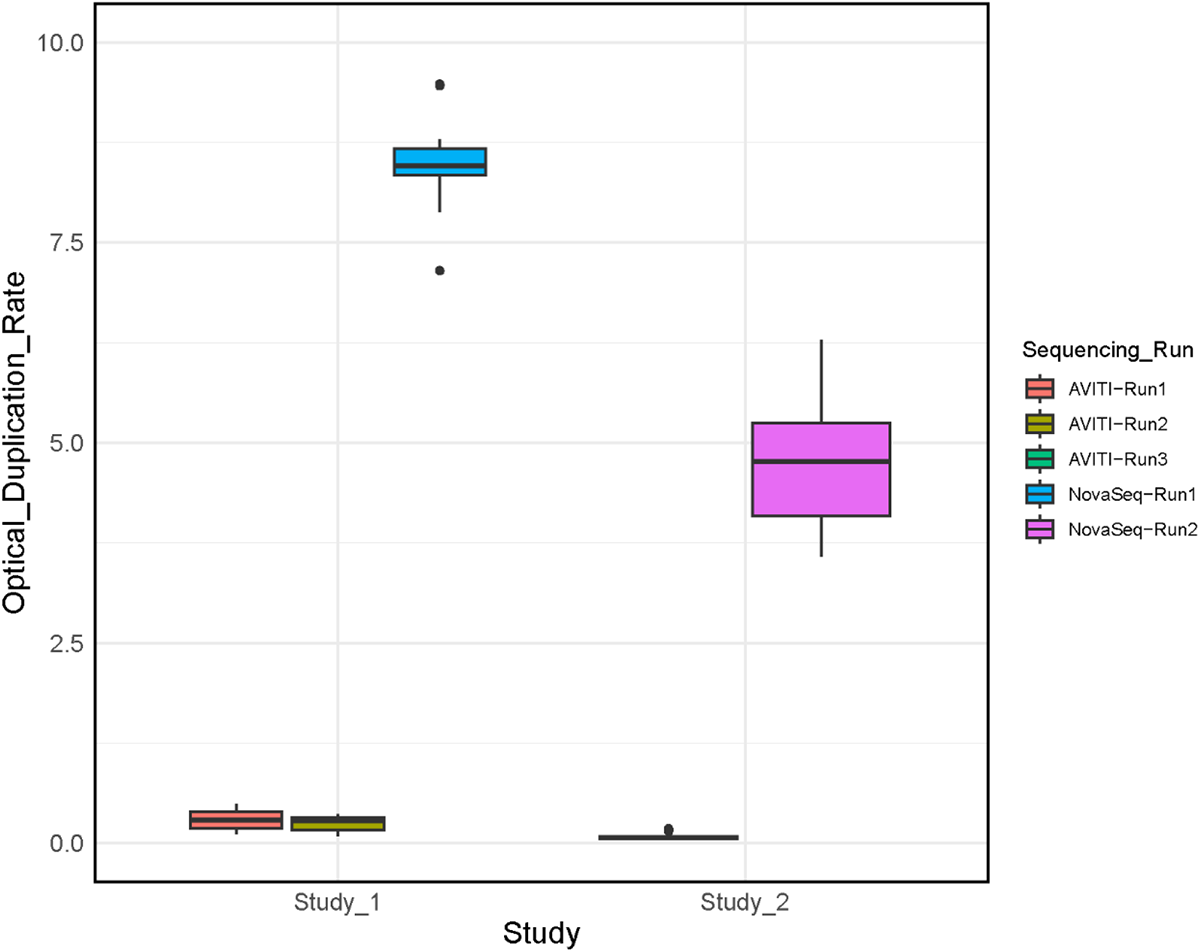
Comparison of duplicate rate between two sequencing platforms X-axis and Y-axis represent study cohort and optical duplicate rate, respectively. Orange, green and purple colors represent sample used for AVITI sequencing. Blue and rose red colors represent samples used for NovaSeq sequencing.

### Quality metrics for assay performance evaluation

We developed a range of sequencing-level and variant calling-level quality metrics to evaluate the sequencing quality for wide range of variant types covered in OncoScreen^TM^ Plus assay. QC data generated from both sequencing systems were compared to evaluate panel performance (Figure 4). As discussed previously, AVITI runs in both Study 1 and 2 showed significantly higher Q30 read percentage than NovaSeq data (Figure 4A). Despite of similar or lower raw reads, AVITI runs in Study 1 and 2 showed comparable or higher mean target coverage due to high data quality and low duplicates percentage (Figure 4B). Although the fold 80 base penalty in AVITI runs was similar to NovaSeq data (Figure 4C), the uniformity was slightly lower than NovaSeq data (Figure 4D). Interestingly, NovaSeq runs from Study 1 and 2 showed higher AT dropout than that from AVITI data (Figure 4E). This GC/AT dropout profile is more similar to NextSeq based on our historical observations. AVITI runs in Study 1 showed higher GC dropout than NovaSeq data (Figure 4F). While in Study 2, the GC profiles were similar between AVITI run 3 and NovaSeq run 2, suggesting sample or library variation could also be a contributing factor.

**Figure 4.**
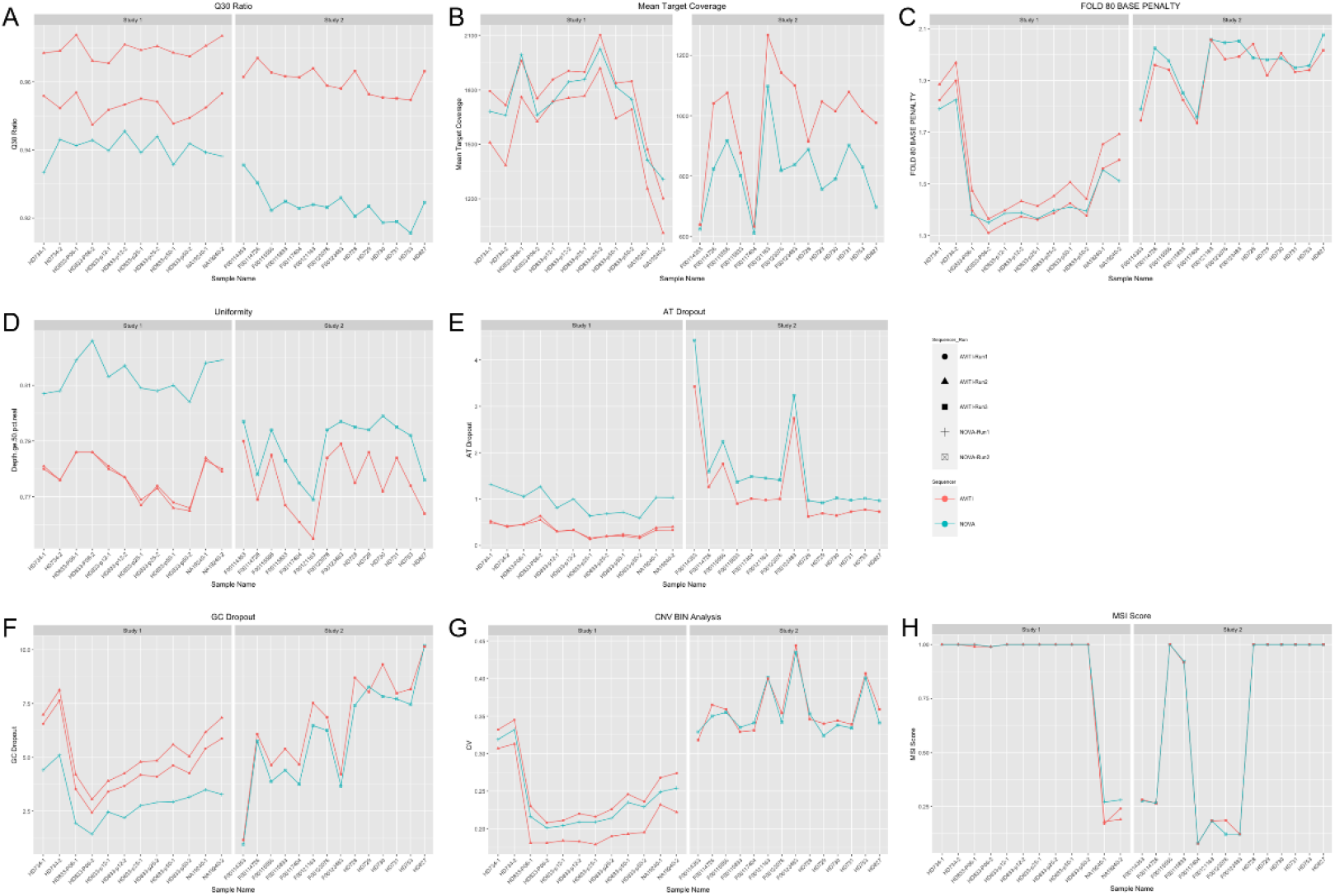
Assay performance evaluation of QC metrics between two sequencing platforms A: Q30 ratio; B: mean target coverage; C: fold 80 base penalty; D: uniformity; E: AT dropout; F: GC dropout; G: CNV BIN Analysis; H: covered loci Num. X-axis and Y-axis represent sample name and QC metrics, respectively. Red and light blue represent AVITI sequencing and NovaSeq sequencing, respectively.

We developed some variant type-specific QC metrics in-house to evaluate the impact of data quality on variant detection success rate. Both systems provided similarly good SNV/Indel variants coverage, precision for CNV detection (Figure 4G), and coverage to identify MSI loci (Figure 4H). To further compare the MSI detection capability of both platforms, the normalized coverage of each MSI site against mean coverage depth was plotted to compare both platforms using NA19240 as the example (Figure 5). The AVITI data showed higher coverage compared to NovaSeq, indicating improved accuracy in short tandem repeats and homopolymers by AVITI. This result substantiates observations made previously in conclusion of the comparison between the AVITI technology and SBS technology in sequencing through long homopolymers (8). Taken as a whole, these results demonstrate that the AVITI system produced sequencing data compatible with the existing pipelines and is suitable for comprehensive genomic profiling applications in oncology. 3732.6+2564.92

**Figure 5.**
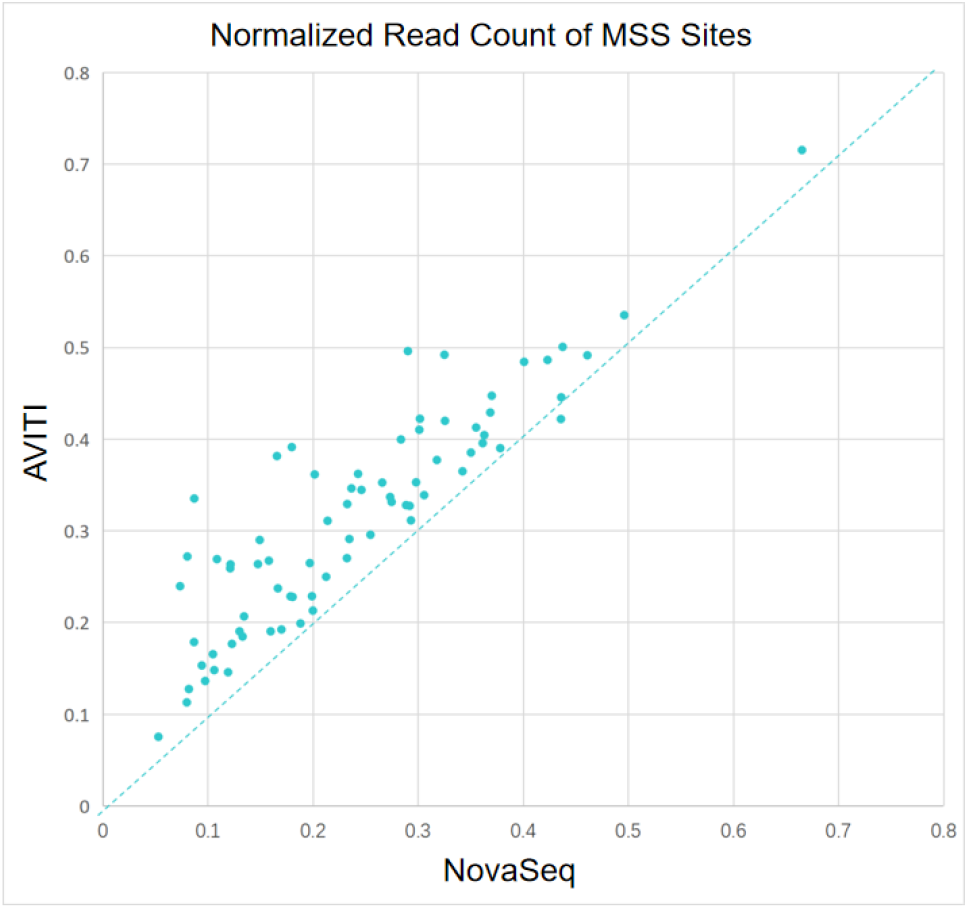
Comparison of detection capability of MSI between two sequencing platforms X-axis and Y-axis represent AVITI sequencing and NovaSeq sequencing, respectively.

### Study 1: Variant detection using reference control samples

Variant detection performance was first investigated using commercial reference control samples (alone or in serial dilution mix). NA19240, HD734 1.3% Tier reference gDNA control, and a set of serial diluted HD833 OncoSpan cfDNA control samples with a “wild-type” background of NA19240 gDNA were included in Study 1. Detection of variants covered by OncoScreen^TM^ Plus panel in a serially diluted HD833 samples are shown in Table 3 and Table 4. Based on reference control material documentation, a total of 72 validated variants are expected to be detected in the OncoScreen ^TM^ Plus target region. It is noted that 3 variants out of 72 were consistently not-detected on both platforms, likely due to their locations in the repeat regions. Limiting the analysis to the remaining 69 annotated variants, greater than 97% of variants were correctly reported from 1/2, 1/4 and 1/8 dilutions in both data sets. Notably, the variant calling pipeline used for this analysis was not designed and trained with AVITI data. At 4,000x coverage, more than 94% of the 69 annotated variants in all dilutions were detected except the 1/16 dilution tier within both AVITI and NovaSeq data sets (Table 3), indicating the limit of detection (LoD) in this assay. The serial diluted HD833 data were leveraged to impute “true” variant calling of 425 variants based on gradual reduction of allele frequency at different dilution levels (Table 4). Among the 425 variants evaluated, greater than 98% were reported via our analysis pipeline in the 1/2 and 1/4 dilutions in both data sets. More than 88% at 1/8 dilution, and more than 56% with 1/16 dilution out of 425 annotated variants were detected in both data sets. Some non-detected variants in 1/8 and 1/16 dilutions are below the LOD for the OncoScreen Plus assay). In addition, allele frequency filter settings may account for the difference in variant detections at low dilution levels. Based on the 425 variants evaluated in Study 1, correlations of variant frequency detected in AVITI data and NovaSeq data were high at higher dilution tiers (R^2^ at 0.97 in 1/2 dilution, 0.95 in 1/4 dilution, and 0.83-0.90 from 1/8 dilution) (Figure 6A-C), and lower with further diluted (0.62-0.71 at the 1/16 dilution sample – to evaluate 1-2% variant frequency near or below LoD of this assay) (Figure 6D). High concordance of both MSI and TMB scores in serial diluted control samples was observed on both sequencing platforms (Figure 6E).

**Figure 6.**
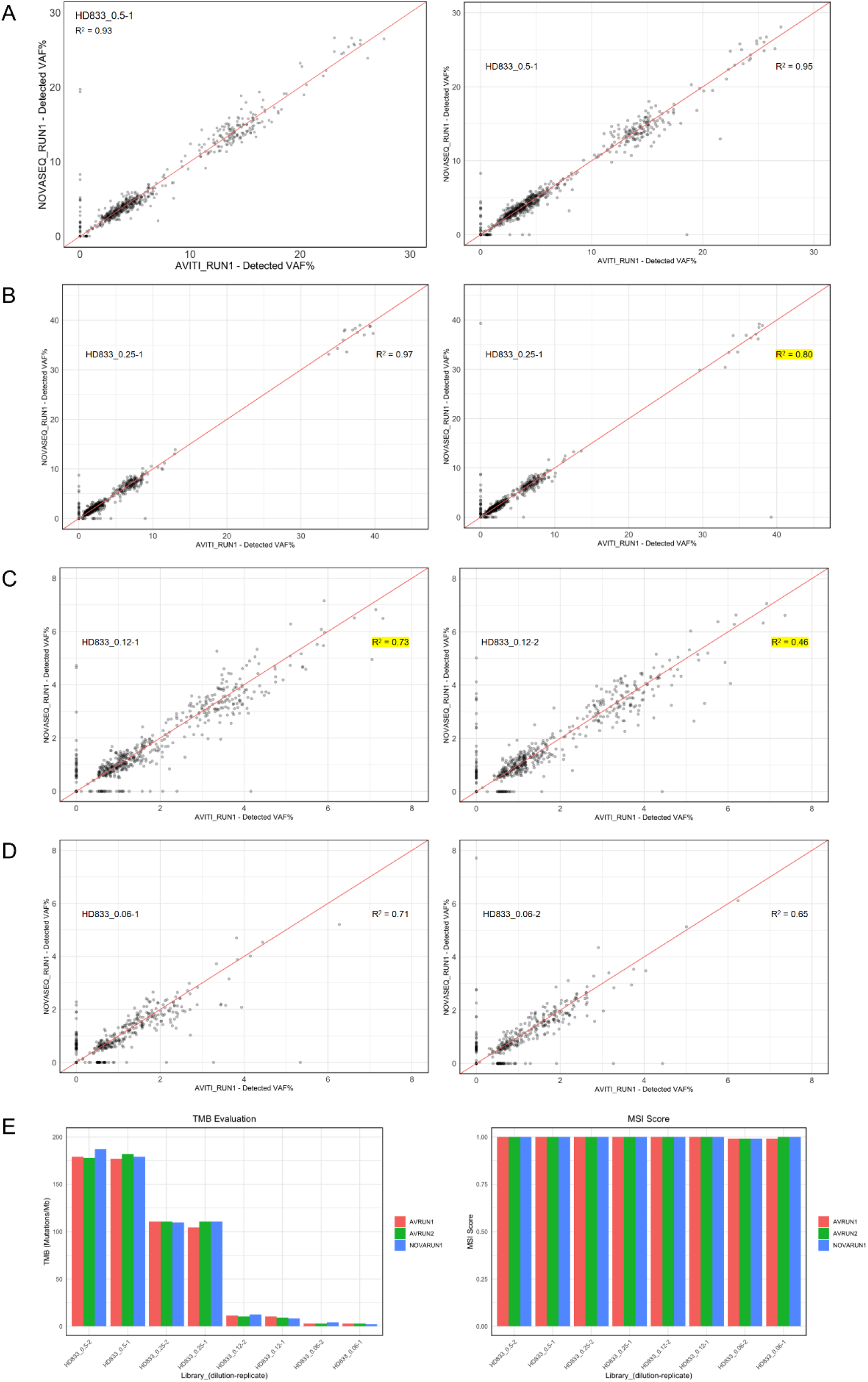
Vairant detection in HD833 series samples in Study 1 between two sequencing Platforms A: correlation analysis of variant frequency in 1/2 diluted sample. X-axis and Y-axis represent run 1 of AVITI sequencing and NovaSeq sequencing, respectively; B: correlation analysis of variant frequency in 1/4 diluted sample. X-axis and Y-axis represent run 1 of AVITI sequencing and NovaSeq sequencing, respectively; C: correlation analysis of variant frequency in 1/8 diluted sample. X-axis and Y-axis represent run 1 of AVITI sequencing and NovaSeq sequencing, respectively; D: correlation analysis of variant frequency in 1/16 diluted sample. X-axis and Y-axis represent run 1 of AVITI sequencing and NovaSeq sequencing, respectively; E: detection of MSI score and TMB evaluation in serial dilution samples. X-axis and Y-axis represent sample and score, respectively. Blue, orange and green represent run 1 of AVITI sequencing, run 2 of AVITI sequencing and NovaSeq sequencing, respectively.

**Table 3.** Detection of variants covered by the OncoScreen Plus panel.

| Sample | Repeat | Based on 72 annotated variants |  | Based on 69 annotated variants |  |
| --- | --- | --- | --- | --- | --- |
|  |  | AVITI | NOVA | AVITI | NOVA |
| **HD833 (1/2)** | Repeat 1 | 94.44% | 94.44% | 98.55% | 98.55% |
|  | Repeat 2 | 94.44% | 94.44% | 98.55% | 98.55% |
| **HD833 (1/4)** | Repeat 1 | 94.44% | 95.83% | 97.10% | 97.10% |
|  | Repeat 2 | 95.83% | 95.83% | 98.55% | 97.10% |
| **HD833 (1/8)** | Repeat 1 | 94.44% | 94.44% | 97.10% | 97.10% |
|  | Repeat 2 | 94.44% | 94.44% | 97.10% | 97.10% |
| **HD833 (1/16)** | Repeat 1 | 87.50% | 87.50% | 91.30% | 91.30% |
|  | Repeat 2 | 86.11% | 86.11% | 89.86% | 89.86% |

**Table 4.** Detection of 425 variants covered by OncoScreen Plus panel.

| **Sample** | **Repeat** | **AVITI** | **NOVA** |
| --- | --- | --- | --- |
| **HD833 (1/2)** | Repeat 1 | 99.06% | 99.29% |
|  | Repeat 2 | 99.53% | 99.06% |
| **HD833 (1/4)** | Repeat 1 | 98.82% | 99.29% |
|  | Repeat 2 | 99.53% | 100.00% |
| **HD833 (1/8)** | Repeat 1 | 89.88% | 89.18% |
|  | Repeat 2 | 88.24% | 88.24% |
| **HD833 (1/16)** | Repeat 1 | 58.82% | 59.76% |
|  | Repeat 2 | 57.65% | 56.94% |

### Variants detection with reference samples in Study 2

Table 5 presents the variant detection performance of “known” SNV/Indels in reference control samples covered by the OncoScreen^TM^ Plus panel. At 4,000x coverage, 100% of documented variants listed for HD728, HD729, HD730, HD731, HD753, HD827 samples were detected in both AVITI and NovaSeq data. For HD734, 5 or 8 variants were not detected by either platform, which is likely due to the variant frequency below LoD (< 2%). Although the variant calling pipeline used for this analysis was not optimized with AVITI data, the analysis result from AVITI data without additional preprocessing or pipeline tuning showed a high concordance of AVITI data in the comparison to NovaSeq data based on all the detected variants of SNV/Indels across the whole targeted region of the OncoScreen^TM^ Plus panel. There were 34 unmatched variants in OncoScreen™ Plus Panel region between NovaSeq and AVITI data in reference sample HD753 (Table 6). 20 of the 34 unmatched variants were due to polish by white blood sample (WBC) background, 8 variants were due to AF filter, 3 variants were due to vertical complex filter, 1 variant was due to not_in_report_region, 1 variant was due to repeat filter, and 1 variant was due to uncall. Potentially due to the wide genome region covered by OncoScreen^TM^ Plus panel instead of the capability differences of the AVITI system and the Nova sequencer, the complexity of the variant detection greatly increased.

**Table 5.** Sensitivity and accuracy of variant detection from reference samples in Study 2.

| **Sample**<br>**name** | **Sequencer** | **Total** | **Detected** | **Undetected** | **PPA** |
| --- | --- | --- | --- | --- | --- |
| **HD728** | AVITI | 13 | 13 | 0 | 100.00% |
|  | NOVA | 13 | 13 | 0 | 100.00% |
| **HD729** | AVITI | 13 | 13 | 0 | 100.00% |
|  | NOVA | 13 | 13 | 0 | 100.00% |
| **HD730** | AVITI | 14 | 14 | 0 | 100.00% |
|  | NOVA | 14 | 14 | 0 | 100.00% |
| **HD731** | AVITI | 12 | 12 | 0 | 100.00% |
|  | NOVA | 12 | 12 | 0 | 100.00% |
| **HD734-1** | AVITI | 43 | 35 | 8 | 81.40% |
|  | NOVA | 43 | 38 | 5 | 88.37% |
| **HD734-2** | AVITI | 43 | 35 | 8 | 81.40% |
|  | NOVA | 43 | 35 | 8 | 81.40% |
| **HD753** | AVITI | 14 | 14 | 0 | 100.00% |
|  | NOVA | 14 | 14 | 0 | 100.00% |
| **HD827** | AVITI | 66 | 66 | 0 | 100% |
|  | NOVA | 66 | 66 | 0 | 100% |

PPA: positive percentage agreement.

**Table 6.** Unmatched variants in OncoScreen™ Plus Panel region between NovaSeq and AVITI data in reference sample HD753.

| ID | Gene | Mutation | Variant<br>Type | AF |  | Filter |
| --- | --- | --- | --- | --- | --- | --- |
|  |  |  |  | AVITI | NOVA |  |
| 1 | ARID1A | p.M1564fs | INDEL | 6.98% | -- | Uncall |
| 2 | GATA2 | p.G200fs | INDEL | 5.18% | -- | polish by wbc bg |
| 3 | EPHA5 | p.R701fs | INDEL | 5.73% | -- | polish by wbc bg |
| 4 | FANCE | p.S154F | SNV | 2.78% | -- | polish by wbc bg |
| 5 | CDK6 | c.834+3A>G | SNV | 2.71% | -- | AF_filter |
| 6 | PRKDC | p.S75N | SNV | 3.39% | -- | polish by wbc bg |
| 7 | CCND1 | p.P287L | SNV | 2.78% | -- | polish by wbc bg |
| 8 | MLH3 | p.K585fs | INDEL | 10.95% | -- | polish by wbc bg |
| 9 | AXIN1 | c.2294+8_2294+30del | INDEL | 4.22% | -- | vertical complex<br>filter |
| 10 | FANCA | p.P615fs | INDEL | 6.61% | -- | polish by wbc bg |
| 11 | HOXB13 | p.G226E | SNV | 3.46% | -- | AF_filter |
| 12 | RNF43 | c.253-8C>G | SNV | 3.96% | -- | polish by wbc bg |
| 13 | PIK3R2 | p.V425L | SNV | 3.79% | -- | polish by wbc bg |
| 14 | AXL | p.P337fs | INDEL | 5.18% | -- | polish by wbc bg |
| 15 | ZNRF3 | p.H705fs | INDEL | 6.68% | -- | polish by wbc bg |
| 16 | AR | p.A52T | SNV | 6.01% | -- | AF_filter |
| 17 | EPHA2 | p.P460fs | INDEL | -- | 5.78% | polish by wbc bg |
| 18 | TENT5C | p.L288S | SNV | -- | 3.43% | AF filter |
| 19 | MSH2 | p.G18= | SNV | -- | 3.15% | AF filter |
| 20 | EPHB1 | p.G628= | SNV | -- | 4.68% | AF filter |
| 21 | CYP17A1 | p.K59fs | INDEL | -- | 5.18% | polish by wbc bg |
| 22 | TCF7L2 | p.P99R | SNV | -- | 4.45% | AF filter |
| 23 | FGFR2 | p.N184fs | INDEL | -- | 5.20% | polish by wbc bg |
| 24 | KMT2A | p.P773fs | INDEL | -- | 10.08% | polish by wbc bg |
| 25 | MLH3 | p.Q341* | SNV | -- | 2.53% | polish by wbc bg |
| 26 | SLX4 | p.G1494_S1508del | INDEL | -- | 5.84% | vertical complex<br>filter |
| 27 | MAPK3 | p.A52= | SNV | -- | 4.98% | not_in_report_region |
| 28 | FANCA | c.3626+4C>T | SNV | -- | 3.46% | AF filter |
| 29 | ALOX12B | p.L451fs | INDEL | -- | 12.93% | polish by wbc bg |
| 30 | NCOR1 | p.T525fs | INDEL | -- | 10.62% | repeat filter |
| 31 | PTPRS | c.2495-11T>C | SNV | -- | 14.55% | vertical complex<br>filter |
| 32 | NOTCH3 | c.5363-17_5363-16inv | COMPLEX | -- | 7.66% | polish by wbc bg |
| 33 | POLD1 | p.D987fs | INDEL | -- | 16.26% | polish by wbc bg |
| 34 | GNAS | p.D167H | SNV | -- | 4.21% | polish by wbc bg |

### Variant detection with clinical FFPE samples in Study 2

Lastly, variant detection performance was evaluated using 8 clinical FFPE samples from a variety of cancer types (endometrial, gastric, pancreatic, colorectal, and head and neck). The frequency of variants detected by both AVITI and NovaSeq data sets showed a high correlation (R^2^ > 0.95 in 7 samples) (Figure 7). Certain variants were not called in one of either the AVITI or NovaSeq data in sample F00115556 (R^2^ at 0.91). FFPE sample quality, variant frequency, or variant complexity of the OncoScreen^TM^ Plus panel are potential causes. The pipeline reported high concordance of CNV, MSI, and TMB scores in 6 of the 8 FFPE sample data sets from both platforms (Figure 9). Overall, data concordance between AVITI and NovaSeq platforms further supports the conclusion that the AVITI system is suitable for sequencing comprehensive genomic profiling assays for oncology applications and provides concordance of detection in a variety of variant types as compared to existing SBS technology.

**Figure 7.**
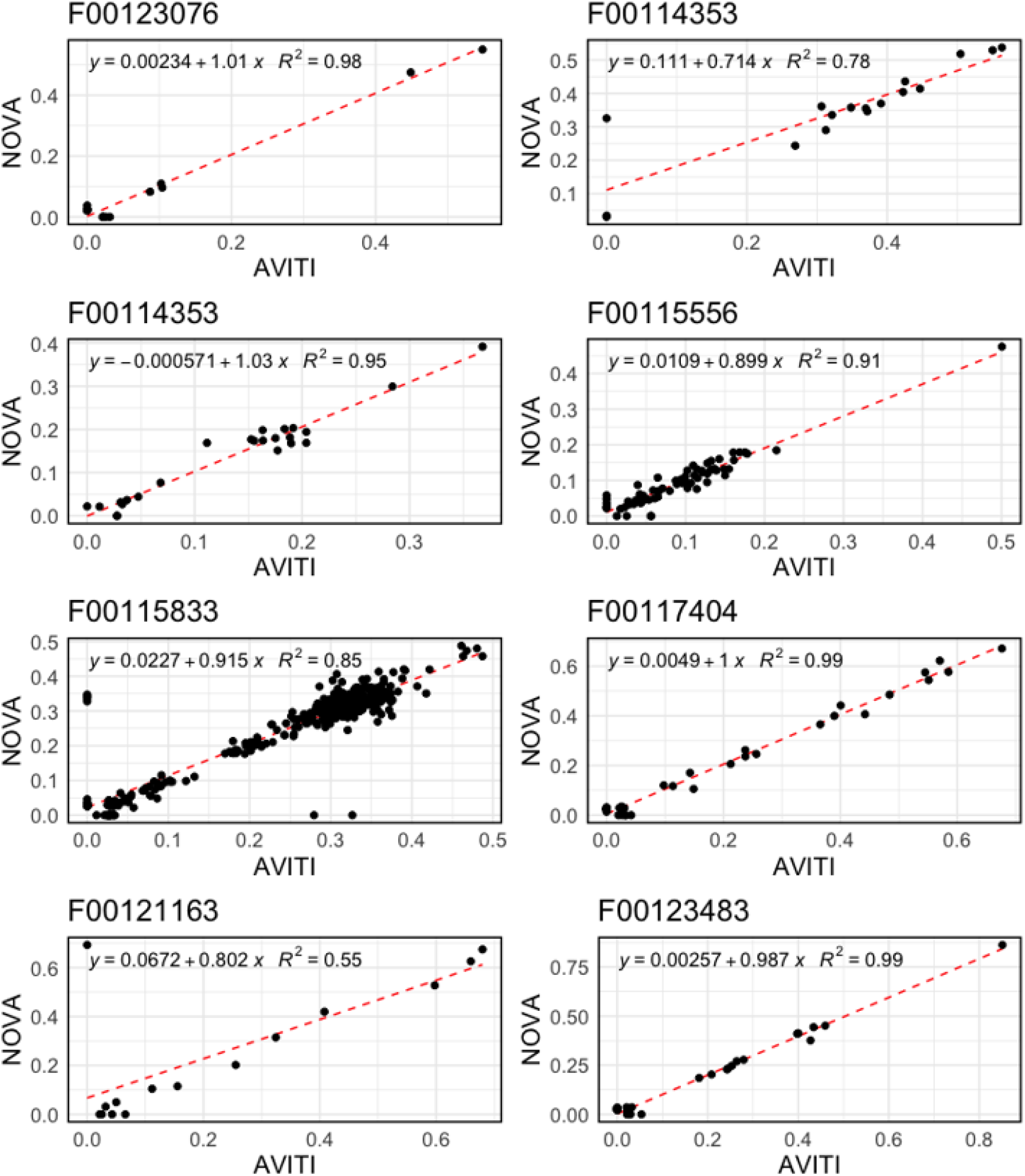
Correlation analysis of variant frequency in clinical FFPE samples in Study 2 between two sequencing platforms X-axis and Y-axis represent AVITI sequencing and NovaSeq sequencing, respectively.

**Figure 8.**
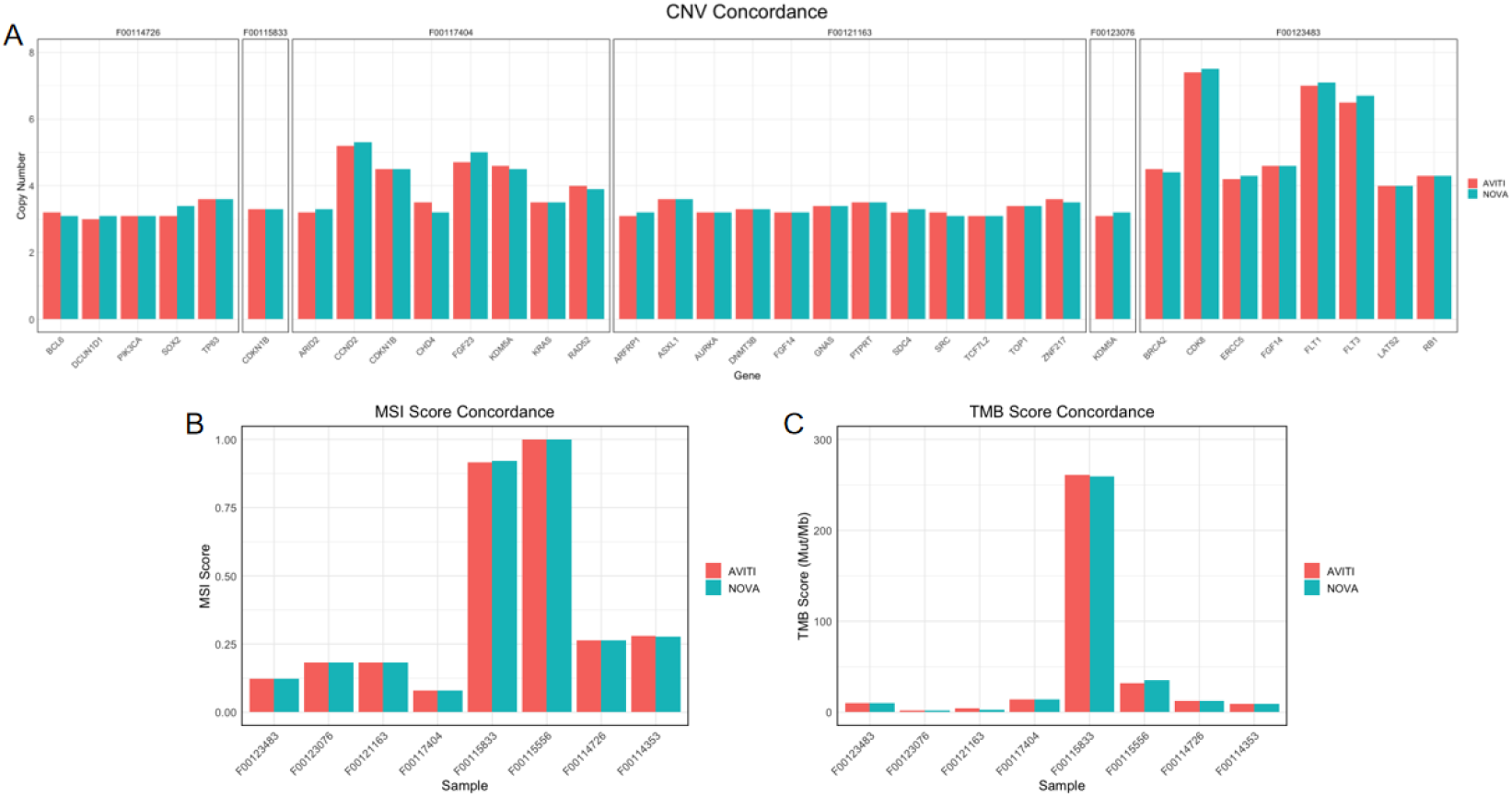
Concordance evaluation of CNV, MSI, and TMB detection in clinical FFPE samples in Study 2 between two sequencing platforms Blue and orange represent AVITI sequencing and NovaSeq sequencing, respectively.

## Discussion

For the last decade, next generation sequencing has been implemented widely in precision oncology and NGS assays have gained regulatory status for companion diagnostics. Target capture with designed Oncology CGP panels has been one of the most cost-effective NGS methods to assess the breadth of cancer related genomic alterations and variant signatures in a single sequencing assay. In addition to sample collection, nucleic acids extraction, and library preparation, high quality sequencing is crucial to provide accurate variant calling and reduce the impact from false positive calls. With the rapid development of new sequencing technologies, it is needed to compare different platforms to maintain an accurate understanding of their relative performance.

In this study, a new commercialized sequencing platform AVITI with a novel sequencing technology, Sequencing by Avidity, was evaluated. Known reference control DNA samples and clinical research FFPE samples were processed through a target enrichment workflow, followed by sequencing on both Element AVITI system and Illumina NovaSeq 6000 platform. Our results showed that AVITI sequencing provides accurate detection of cancer-related variants and is compatible with the OncoScreen^TM^ Plus assay with existing variant calling analysis pipelines. This compatibility may enable users to test the sequencing system and generate high quality sequencing data with minimum transition cost without the requirements of batching high sample quantity in order to meet the cost-effective sequencing run provided by NovaSeq system.

Higher percentage of quality score and mean target coverage, and lower optical duplication rate was observed on AVITI platform, compared with NovaSeq sequencing platform. Sequencing quality scores can predict actual error rates and identify high-quality bases (12). A higher quality score indicates a lower probability of an error. Low optical duplication rate in sequencing contributes to better data diversity and efficiency. The lower optical duplication rates (∼ 1% on AVITI vs ∼ 8% on NovaSeq) and high quality of the sequencing runs from AVITI resulted in more usable reads (99% of output reads on AVITI vs ∼ 92% of output reads on NovaSeq), indicating that lower data output is required to reach similar targeted coverage on the AVITI platform.

Data generated from AVITI system were used directly as input in the variant calling pipeline and compare a full spectrum of variant types in CGP assay, including SNVs, Indels, CNV, MSI, and TMB. Both AVITI data and Illumina NovaSeq data identified the vast majority of variants and showed high concordance of variant allele frequencies between the two platforms (> 90% in 5 out of 8 FFPE samples) and comparing to known reference was high (> 95%) except for HD734 reference control harboring several variants with AF lower than LoD. This result suggests that the “Sequencing by Avidity” technology is suitable for somatic variant detection in identifying variant frequency at the comparable level around 2% with 4000x coverage and 30 ng gDNA input as SBS technology. It is also worth mentioning that AVITI showed better MSI coverage due to better homopolymer sequencing capabilities.

In studies with serial dilution of reference control samples, a small subset of variants fell below the limit of detection of the OncoScreen Plus assay (0.5%), which were not reported in data from both platforms. Some variants were called in one platform but not the other. The investigation of this difference revealed several factors. Firstly, coverage uniformity of the same sample was slightly different on two platforms. With higher GC dropout from AVITI data and higher AT dropout from NovaSeq data, might indicate the amplification mechanism could play a role in favoring GC-or AT-rich content. The overall lower uniformity from AVITI data suggested that sequencing conditions can be further improved, and panels and analysis parameters optimized on one platform could show slightly different performance on other platforms, although in this test case, good coverage and high concordance were observed despite of uniformity difference. Another factor is the library conversion step. To sequence libraries made for Illumina sequencers, circularization of the library pool is required for loading on AVITI system. Amplification of circular library in AVITI system utilizes the original library molecule as template for amplification, resulting in the advantage of minimizing the magnification of potential amplification errors and better accuracy in sequencing (8). Whether this circularization process may introduce variability or possible different bias is worth further investigation. Although the diversity and representation are highly comparable between platforms in this study, additional studies will be useful with other assay types or additional experiments to show broader reproducibility. Finally, input sample type and library preparation are also important factors to consider in comparative experimental design. These are potential contributing factors in the observation that GC bias and sequence uniformity is improved on the AVITI platform moving from Study 1 to Study 2.

Another potential source of differences in variant detection in this comparison study is the pipeline used for analysis. Data generated from AVITI system in this study were directly used in the analysis pipeline, which was designed based on historical data from Illumina sequencing and quality score system. The analysis pipelines mainly consist of read alignment, quality assessment, variant identification and annotation. Different combinations of tools will lead to the divergence of performance among pipelines and finally affect the final interpretation of variants calling results (13). The most accurate identification of genomic variants requires standardization of benchmarking performance and optimization of analysis pipelines for genomics research based on NGS technology. Focused optimization and algorithm training for variant calling algorithm based on Element AVITI system may further improve the variant detection accuracy.

Reliable and proper reference DNA is critical in variant detection panel development and assay evaluation. In this study, a variety of commercially available control reference genome samples were compared. Often, more variants were reported than those listed by the vendors. Data generated from AVITI may serve as a benchmark reference providing high coverage of targeted regions in samples with a given assay. In addition, our control sample titration design was further examined to define a more comprehensive variant list in HD833. The list of variants covered in our OncoScreen^TM^ Plus panel is provided as a supplementary document (supplementary Table 1). Researchers using the OncoScreen Plus assay or other panels covering similar regions may use this dataset to verify variants and fine tune variant calling filter parameter settings.

In conclusion, this study is the first to compare sequencing performance on Oncology CGP panels between the AVITI system and Illumina NovaSeq 6000 platforms. Both library preparation workflow and output data format from the AVITI system were fully compatible with the OncoScreen^TM^ Plus assay and associated analysis pipeline. Moreover, both the AVITI and Illumina NovaSeq platforms provided high concordance rates in the detection across somatic variant types with reference genome and cancer FFPE samples. Data generated from the AVITI system can be used to compare continuous study in projects that might have samples sequenced on different platforms.

